# Light-regulated pro-angiogenic engineered living materials

**DOI:** 10.1101/2022.10.28.514190

**Authors:** Priyanka Dhakane, Varun Sai Tadimarri, Shrikrishnan Sankaran

## Abstract

Despite their promise, the application of growth factors in regenerative medicine is limited by their poor stability in the body, high costs of production/storage and need for localized and tightly controlled delivery to minimize adverse side effects. In this study, a unique strategy to overcome these limitations is explored based on engineered living materials (ELMs). These are an emerging class of composite materials, which contain live microorganisms that can be engineered to produce and secrete proteins in response to external stimuli. Herein, the development of an ELM that light-responsively releases a pro-angiogenic protein is described. This is achieved by optogenetically engineering bacteria to synthesize and secrete a fusion protein containing a vascular endothelial growth factor peptidomimetic linked to a collagen-binding domain. The bacteria are securely encapsulated in bilayer hydrogel constructs that support bacterial functionality but prevent their escape from the ELM. The possibility to switch protein release ON and OFF with light and to tune the amount released with different light intensities is demonstrated. Finally, it is shown that the released protein is active through its ability to bind to collagen and promote angiogenic network formation in human vascular endothelial cell cultures, indicating the regenerative potential of these ELMs.

## II. Introduction

Regenerative medicine is a rapidly developing research field focused on accelerating the repair of damaged cells, tissues and organs to restore normal function and circumvent the need for translplantation^1^. In healing processes, growth factors play a major role in stimulating cells and orchestrate transformations in them necessary for regeneration. They are proteins that are secreted by cells, bind to the surrounding extracellular matrix (ECM) and interact with the receptors on the surfaces of other cells. These receptor-specific interactions trigger signal transduction pathways that promote events such as cell growth, proliferation, differentiation, cell migration, adhesion and survival^2–4^. For instance, vascular endothelial growth factor (VEGF), and platelet derived growth factor (PDGF) strongly stimulate blood vessel formation increasing the permeability of endothelial cells and by activating integrin, NOTCH and WNT signalling for endothelial cell differentiation.^5^ Thus, growth factors are powerful tools in regenerative medicine. However, due to their potency in driving cellular transformations, their concentration and localization need to be carefully regulated. Failing to do so can cause overstimulation or off-site differentiation of cells, leading to necrosis or tumorigenesis.^6,7^ This is exemplified by the long list of severe side-effects plaguing the few clinically approved growth factor-based medical products like PDGF-based Regnarex (side effects include increased risk of systemic cancer, skin rash and cellulitis)^8^. Apart from unwanted side effects, growth factors are also complex bulky proteins often with poor stability, making them expensive to produce, purify, store, and deliver at effective doses^7,9^ These technical challenges have thus far limited the clinical applicability of growth factor-based therapies despite over 3 decades of research demonstrating their potential.^10^

These issues are being addressed through 2 major strategies – (i) use of drug release systems ensuring localized and controlled supply of the growth factor over time and (ii) development of short and robust variants mimicking a desired function of the growth factors. This is particularly exemplified by advances in VEGF-based therapies. This growth factor plays a prominent role in stimulating the sprouting of new blood vessels from existing ones to supply oxygen to tissues suffering from hypoxia.^11,12^ VEGF based therapies are being explored for peripheral vascular disease (PVD) that results in severe blockage of arteries of the lower extremities, has a high limb amputation and mortality rate and generally has poor prognosis.^13^ In animal models of PVD, increasing VEGF levels to enhance collateral flow around blocked blood vessel has been achieved by intramuscular injection and vascular infusion of an adenoviral vector encoding VEGF^14^. Unfortunately, overexpression or overstimulation by VEGF in laboratory animals can lead to a variety of side-effects like formation of leaky vessels, metabolic dysfunction, transient edema^15,16^, increase in atherosclerotic plaques or uncontrolled neoangiongenesis (increased risk of cancer). Controlled release of VEGF through implantable devices is challenged by this protein’s complexity and low serum stability, leading to high costs, low packing densities and poor control over its release from implantable drug-release devices.^17^ Thus, despite its promise, clinical application of VEGF for PVD has been severely limited by these issues^18,19^.

A cheaper, shorter and more robust alternative to VEGF under investigation is the peptidomimetic, QK (KLTWQELYQLKYKGI), that has been shown to promote formation and organization of capillaries both *in vitro*^20^ and *in vivo*^21^. The peptide has higher potency to drive angiogenic morphogenesis in endothelial cells when immobilized on hydrogel matrices compared to its soluble form.^20,22^ This mimics the activity of VEGF when immobilized to heparan sulphate in the ECM. Furthermore, such immobilization ensures that the pro-angiogenic activity is spatially confined. Recently, sustained release of QK from bone graft materials for up to 6 days was shown to induce angiogenic differentiation in vascular endothelial cells.^23^ Thus, QK has shown great promise for promoting blood vessel formation similar to VEGF. For effective treatment of PVD with wide variabilities in disease profiles and progression among patients, it is desirable to develop a versatile strategy to deliver QK where and when it is required in a cost-effective manner.

In the past 5 years, a unique strategy to achieve low-cost and in situ controllable drug release has been explored in the form of engineered living materials (ELMs). In ELMs, living cells are combined with non-living materials to create composites with programmable and life-like capabilities.^24^ ELMs for drug delivery have been made encapsulating within hydrogels, bacteria genetically engineered to produce and release drugs on demand, remotely controlled by external stimuli like chemical inducers or light. The bacteria in these ELMs can thrive on nutrients available at the disease site and can be triggered to produce and secrete drugs at desired doses when needed. Using hydrogels made of agarose, Pluronic F127 or collagen and bacteria like *E. coli, L. lactis* and *B. subtilis*, therapeutic ELMs have been made in the form of discs, films, patches and 3D printed structures to suit different therapeutic needs. While most studies report the release of anti-microbial drugs from ELMs, one set of studies from the Salmeron-Sanchez lab demonstrates the release of BMP, controlled by a peptide-inducer, nisin.

In this study, an ELM capable of in situ tuneable release of a collagen-binding QK fusion protein is described. For the first time, we demonstrate light controlled release of an active growth factor peptidomimetic from ELMs fabricated in a secure bilayer encapsulation format that prevents bacterial escape/outgrowth.^25^ We show that QK secretion can be sustained in a physiologically relevant range of concentrations (5 – 25 nM) for at least 9 days in a light-controlled manner and that it can induce angiogenic differentiation in endothelial cells.

## III. Results and Discussions

### A. Light-responsive production, secretion and bioactivity of QK

For this study, *Clearcoli*, an endotoxin-free variant of *E. coli*, was engineered to light-responsively synthesize and secrete a QK-bearing fusion protein (YCQ) as shown in **Figure 1A**. This YCQ fusion protein contained (i) YebF, a carrier protein aiding secretion from *E. coli*,^26,27^ (ii) a collagen-binding domain (CBD, sequence - WREPSFMVLS)^28^ to facilitate immobilization of QK to the extracellular matrix, (iii) a Strep-TagII peptide (WSHPQFEK) for purification and staining^29^ and (iv) the QK peptide. Notably, this design allowed for YebF to be at the N-terminal and QK at the C-terminal, which is necessary for their functionality. As a negative control, a similar fusion protein, YCx, was designed bearing a scrambled-QK peptide (GLKEQSPRKHRLG) at the C-terminal, previously shown to be inactive.^30^ The gene corresponding to the 18.9 kDa YCQ or YCx protein was inserted into the multiple cloning site of an optogenetic plasmid, pDawn,^31^ to achieve light responsive production and secretion. This was confirmed through SDS-PAGE by running the extracellular and cellular fractions from bacterial cultures grown either in white light or in the dark (Figure 1B). In the light exposed cultures, a band around 15 kDa was observed in both the cellular and extracellular fractions as well as after purification. This lower molecular weight was expected as a consequence of cleavage of the 2.2 kDa N-terminal signal peptide of YebF during its secretion into the periplasm through the sec pathway. Western Blot analysis confirmed that this band contained the StrepII-tag (**Supporting information, Figure S1**). MALDI-TOF mass-spec analysis of the protein purified from the cellular fraction revealed a clear peak at 16.7 kDa confirming cleavage of the signal peptide suggesting that most of the intracellular protein resided in the periplasm (**Supporting information Figure S1**). These results confirmed that YCQ can be light-responsively produced and secreted from Clearcoli. We proceeded to test the capability of the protein to bind collagen and trigger angiogenic differentiation in vascular endothelial cells. For these assays, we used a collagen-gelatin mixture (Col-Gel) in a 3:1 ratio by weight to form a thin film gel that has been reported to mimic the extracellular matrix in wounds where collagen is often degraded to remodel the tissue.^32^ Binding of YCQ to collagen was verified by incubating the protein on a photo-patterned substrate of Col-Gel, after which it was immunostained using a primary antibody specific to YebF. Both purified and secreted YCQ were observed to preferentially adhere to Col-Gel patterned regions (**Figure 1C, Supporting information Figure S2**).

**Figure 1.**
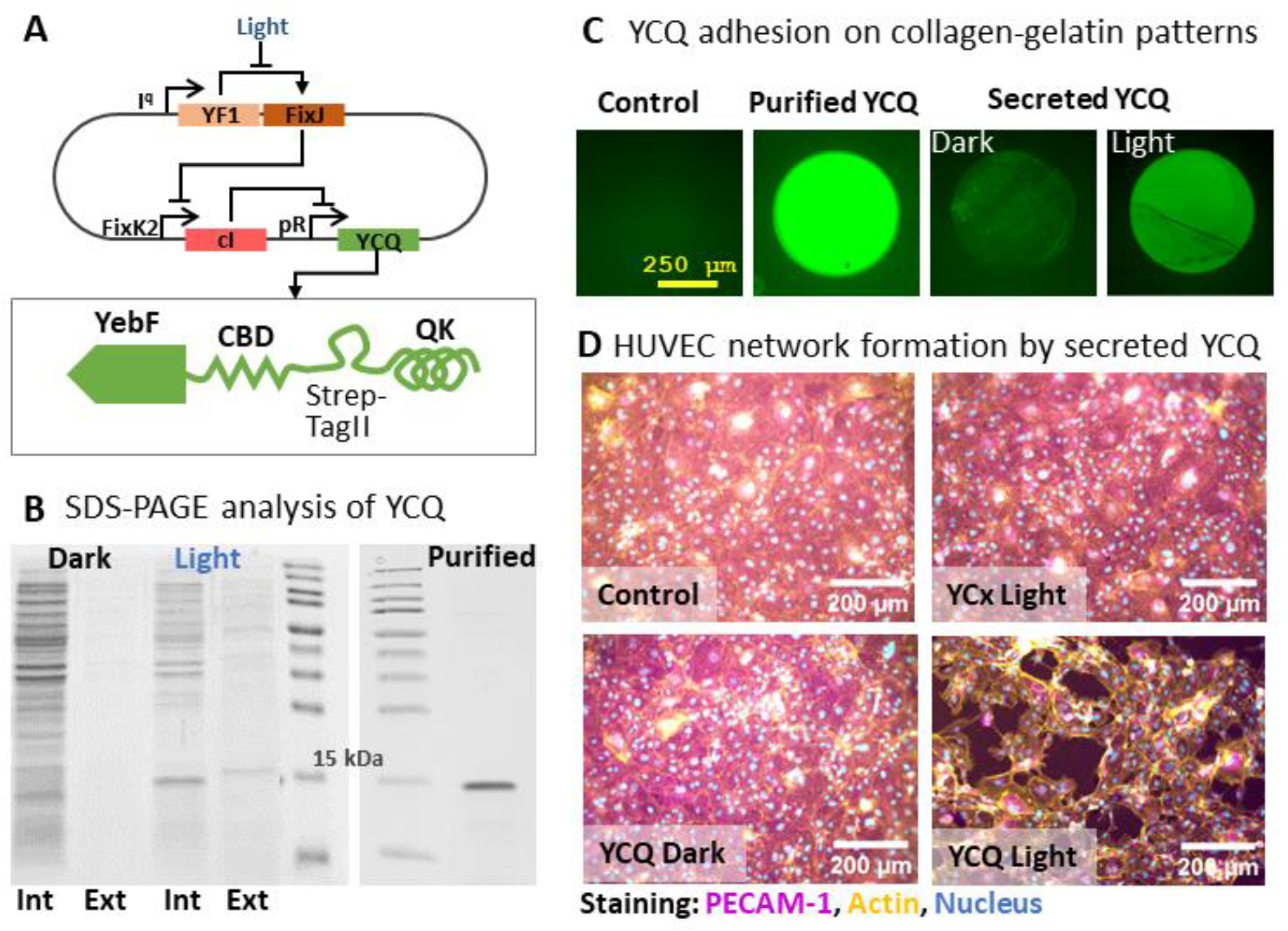
Design and activity of YCQ: A. Graphical representation of the pDawn-based optogenetic circuit for light-responsive production of YCQ along with scheme highlighting the different domains of the fusion protein. B. SDS-PAGE images of the intracellular (Int) and extracellular (Ext) fractions of light-regulated production and secretion of YCQ from engineered bacterial cultures, along with purified YCQ for comparison. C. Fluorescence microscopy images of the Col-Gel photopatterning assay to verify YCQ’s ability to adhere to collagen. Staining was done with an Anti-YebF primary antibody and a fluorescently labelled anti rabbit AF-488 secondary antibody. In the Control condition, the patterned Col-Gel surfaces were incubated with blank medium. D. Fluorescence microscopy images of HUVECs grown for 16 h on Col-Gel surfaces incubated with supernatants of different bacterial cultures grown with or without light. The cells were stained for PECAM-1 (magenta), actin (yellow) and DNA (cyan).

Next, we tested the ability of YCQ-bound Col-Gel matrices to induce angiogenic differentiation in Human Umbilical Vein Endothelial Cell (HUVEC) cultures using a network formation assay.^32,33^Angiogenesis is a vital process for normal tissue development and wound healing, but is also associated with a variety of pathological conditions. Using this protocol, angiogenesis may be measured in vitro in a fast, quantifiable manner. Primary or immortalized endothelial cells are mixed with conditioned media and plated on basement membrane matrix. The endothelial cells form capillary like structures in response to angiogenic signals found in conditioned media. The tube formation occurs quickly with endothelial cells beginning to align themselves within 1 hr and lumen-containing tubules beginning to appear within 2 hr. In summary, this assay can be used to identify genes and pathways that are involved in the promotion or inhibition of angiogenesis in a rapid, reproducible, and quantitative manner^28^For this, cultures of bacteria containing pDawn-YCQ, pDawn-YCx and no plasmid (control) were grown in dark until they reached OD_600nm_ of 0.5 then under light or dark conditions for 12 hours. Cell-free supernatants (SN) from these cultures were then collected, filter sterilized and incubated on Col-Gel surfaces for the proteins to bind with the matrix. As a positive control, 10 nM of purified YCQ was also similarly incubated on the Col-Gel surfaces. When HUVECs were seeded on these Col-Gel surfaces, it was observed that those modified with light-activated YCQ supernatants and purified YCQ promoted the cells to undergo network formation in 16 h (**Figure 1D, Supporting information Figure S3, S4**). On the other hand, on Col-Gel surfaces incubated with control, YCx dark, YCx light and YCQ dark supernatants cells retained monolayer cobblestone morphology over the same period. Immunostaining revealed upregulation of PECAM-1 and actin in cells that formed networks compared to the those that formed a monolayer. These features correlate with the initiation of the angiogenesis process, in line with previous reports. All together, these results confirmed that YCQ performed the secretion, collagen-binding and angiogenesis functions it was designed for.

### B. Engineered living material design and fabrication

To fabricate our ELMs, we encapsulated the bacteria in acrylate modified Pluronic F127 hydrogels, commonly used in medically relevant ELM reports.^25,34^ ensure bacterial survival but prevent their escape to the surroundings, 30% w/v Pluronic F127-diacrylate (PluDA) hydrogels were fabricated in a bilayer format, with bacteria encapsulated in a core layer and surrounded by a protective shell layer. To make the ELMs compatible for microscopy imaging and biochemical assays, they were made in the form of discs bonded to acryloxypropyl silane coated glass cover slips. These constructs were fabricated in a stepwise manner, wherein the bacterial gel was first formed on the glass then coated with the shell hydrogel (**Figure 2A, B**). Chemical cross-linking of the acrylate groups in the gels was done using a photo-initiator (Irgacure 2959) that could be activated using 365 nm light, which is orthogonal to the blue light required to activate YCQ production in the bacteria. The duration (1 min) and power (6 mW cm-2) of the 365 nm irradiation was selected based on a previous report ^25^that identified conditions, which ensured complete cross-linking, while minimally affecting the bacteria. To ensure that the ELMs had well-defined and reproducible dimensions, ring-shaped PDMS moulds (Supplementary information, Figure S5) were used to form the core (dia 6 mm, h 0.8 mm) and shell (dia 13 mm, h 1.2 mm). In this format, the bacterial gel has a volume of almost 23 μL with an initial bacterial density of 0.1 OD_600nm_, resulting in an initial population of ^~^10^6^ bacterial cells (1 OD_600nm_ = 8 × 10^8^ cfu/mL for *E. coli*). When incubated in HUVEC-compliant M199 medium, the bacteria were found to not grow within the gels. To support bacterial growth, the glucose concentration in the medium was increased from 0.1 to 0.5 % w/v and sodium chloride from 0.68 to 1.56 % w/v. HUVEC growth and morphology was not visibly affected by these modifications to the medium (Supplementary information Figure S4). Incubation of the ELMs in the medium at 37 °C resulted in the entrapped bacteria growing from single cells to spherical colonies in 24 h (Figure 2C) after which considerable increase in colony sizes was not observed. No leakage or outgrowth of the bacteria into the medium was observed for at least 15 days, verified by brightfield microscopy (data not shown). Bacterial colonies producing YCQ were stained with fluorescently labelled streptavidin and could be observed as diffuse patches by fluorescence microscopy, possibly due to staining of high concentrations of secreted YCQ in the bacterial colonies (Figure 2C). In comparison, no such patches were observed in gels containing unmodified bacteria (Control).

**Figure 2.**
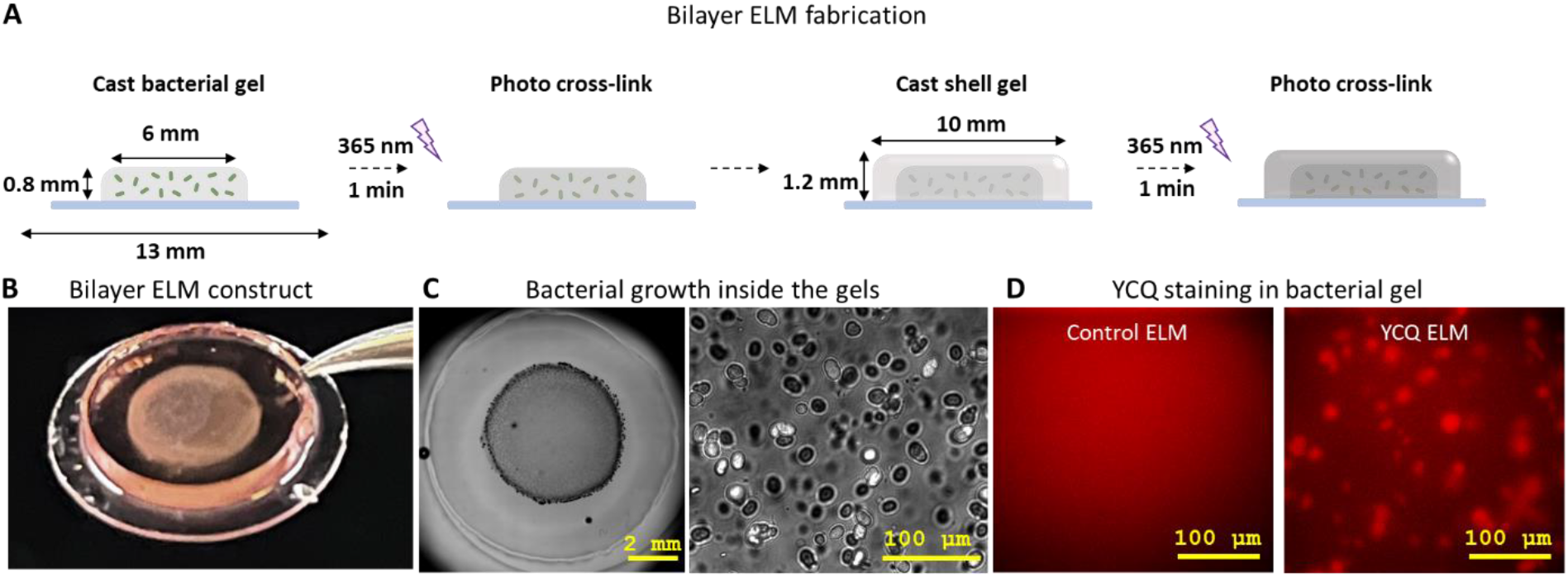
Engineered living material design and fabrication. A. Graphical representation of manual fabrication of ELM. The scheme depicts the steps involved in the making of the bilayer ELMs, along with relevant dimensions of the core and shell layers. B. Macroscopic image of the ELM held by a tweezer. C. Brightfield microscopy stitched image of the ELM with bacteria grown in the core layer (left) along with a magnified image of 6-day grown bacterial colonies. D. Qdot streptavidin staining of YCQ-producing ELM. ClearColi YCQ colonies are stained red indicating YCQ production within ELM; control ELM has unmodified ClearColi.

### C. Light-regulated release of active YCQ from ELMs

Next, we tested the capability of switching and tuning YCQ release from the ELMs using light. After preparation, the ELMs were incubated in dark at 28°C for 16 h, allowing the bacteria to grow into colonies before inducing them with light. Induction of protein expression was maintained by pulsed irradiation with blue light (2 s on, 1 min off, 450nm wavelength, 200uW/cm^2^ power) (**Figure 3A**). The release of YCQ from these bilayer ELM constructs was quantified using a sandwich ELISA assay with a Streptactin-coated plate and a primary antibody specific to YebF (Supporting information, Figure S5). To test the possibility to switch YCQ release ON and OFF, one set of constructs were exposed to blue light for 3 days and another set was left in the dark for the same time at 37 °C (**Figure 3B**). On day 3, YCQ quantification revealed that the amount released in light (^~^1.4 nM) was only slightly higher than what was released in the dark (^~^0.6 nM). The low level of YCQ release in light might be due to bacterial growth continuing to occur in the gel during this period or partial trapping of the protein in the gel. After this, the ELMs that had been exposed to light pulses were placed in the dark and vice versa for another 3 days. At this point, a major difference in production levels were seen between ELMs exposed to light pulses (^~^4 nM) compared to those left in the dark (^~^0.7 nM). Once again, the irradiation conditions were switched for these ELM samples for another 3 days and it was clearly seen that those exposed to light pulses released considerably more YCQ in light (^~^3.8 nM) compared to those in dark (^~^1.1 nM). Furthermore, these results clearly demonstrated that the ELMs could be switched from OFF to ON or ON to OFF state using light. Next, to test the possibility of tuning the amount of YCQ release by varying light intensities, ELMs were induced with pulsed blue light (2s on, 1 min off) with intensities of 0 μw/cm^2^, 80 μW/cm^2^, 105 μW/cm^2^ or 125 μW/cm^2^ for 6 days (**Figure 3C**). The media surrounding ELMs was collected on day 6 from Control (unmodified bacteria) and YCQ-releasing ELMs and was analysed with ELISA (Materials and Methods Section: E, Figure S4). ELMs kept in dark (0 μw/cm^2^) released <1 nM YCQ, those exposed to 80 μW/cm^2^ released 8-15 nM, 105 μW/cm^2^ released 14-19 nM and 125 μW/cm^2^ released 11-20nM of YCQ (figure 3C). These results are suggestive of tuneable control over protein production in ELMs using varying intensities of light within a range of 1 – 20 nM, although smaller deviations under each condition are desirable.

**Figure 3.**
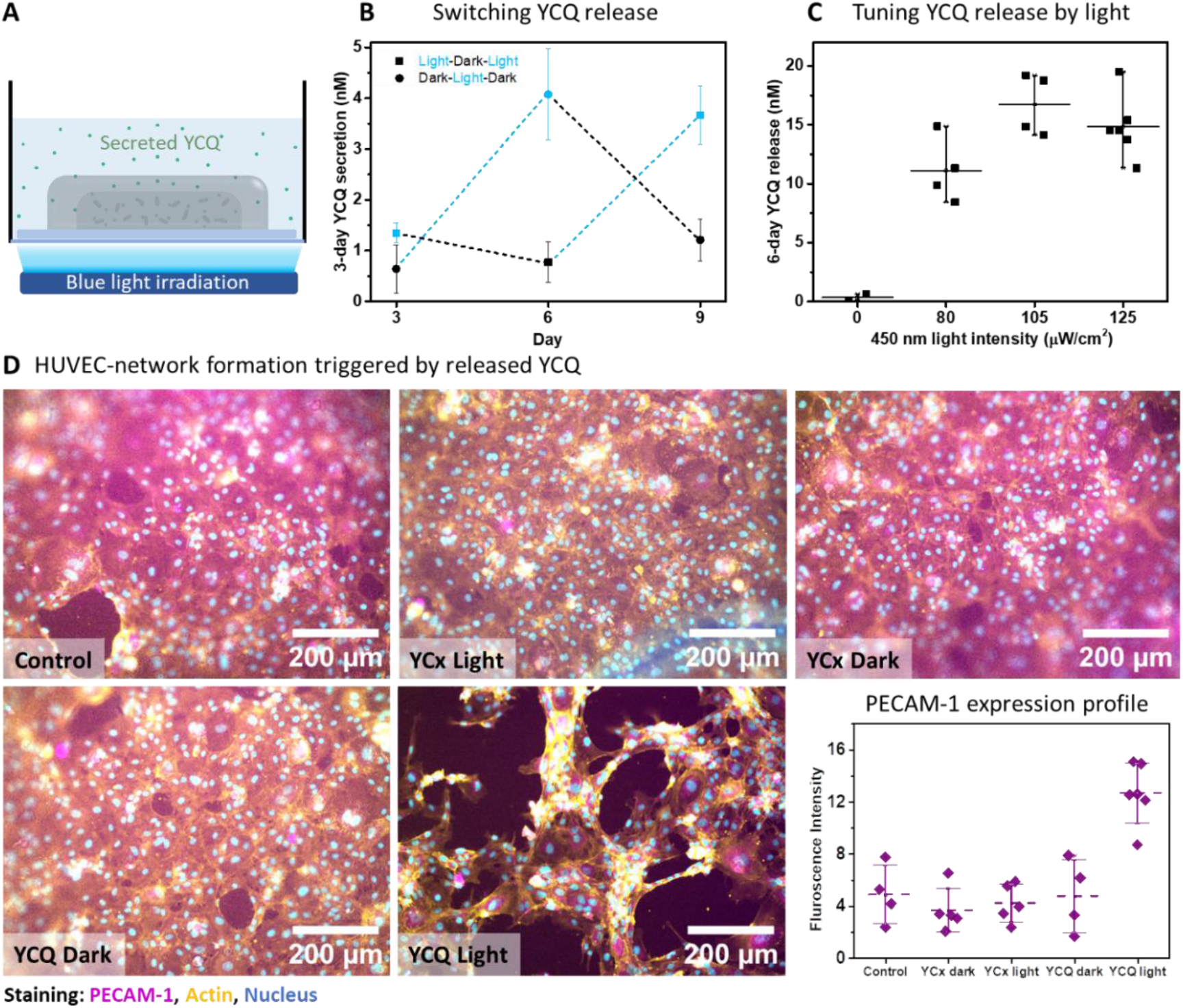
Light regulation of YCQ release from ELMs: A. Graphical representation of the ELM setup including blue light irradiation from below. The ELM constructs were placed in 24 well-plate wells containing 300uL of optimized M199 medium on top of an optowell device for pulsed irradiation at 450 nm (2sec On, 1 min OFF) B. Light switchable control over YCQ release from ELMs. The lines in blue represent durations when samples were placed in light and the lines in black represent durations when samples were placed in the dark. The first blue and black symbols at 3 days were placed in light and dark, respectively till the measurement was made. Symbols represent means values from individual ELM samples and whiskers represent standard deviation C. Tuneable release of YCQ by varying intensities of pulsed blue light irradiation from ELMs. Intensities of 0μW/cm^2^, 80 μW/cm^2^, 105 μW/cm^2^ and 125 μW/cm^2^ were used. Symbols represent individual ELM samples; horizontal bars represent means and whiskers represent standard deviation. D. Epifluorescence images of HUVECs cultured for 16 hours on Col-Gel surfaces that were incubated in supernatants from ELMs containing unmodified bacteria (control), YCx (negative control), and YCQ kept in dark or exposed to blue light pulses (2sec ON, 1min OFF; 105 μW/cm2) for 6 days. The cells were stained for PECAM-1 (magenta), actin (yellow) and DNA (cyan). The graph represents intracellular PECAM-1 signal intensity after 16 hours of seeding HUVECs. Symbols represent mean intensity values from individual images, dashed lines are the means of all data points and whiskers represent standard deviation.

### D. Angiogenesis promoting capabilities of light-responsively secreted YCQ

After establishing the switchable and tuneable release of YCQ from ELMs, we tested the activity of YCQ released from the ELMs after 6 days of pulsed blue light irradiation at 105 μw/cm^2^. As controls, ELMs containing unmodified bacteria and YCx-producing bacteria were also made. The supernatants from the ELMs were incubated with Col-Gel surfaces after which the network formation assay using HUVECs was performed. On 3 out of 4 Col-Gel surfaces incubated with light-exposed YCQ-releasing ELMs, HUVECs were seen to form networks in 16 h, while this did not occur in all other conditions, where HUVECs were seen to grow as monolayers with cobblestone morphology (**Figure 3D, Supporting information Figures S6 – S8**). Notably, this cobblestone morphology was also seen on Col-Gel surfaces incubated with dark-state YCQ-releasing ELMs, indicating that the light-induced fold change of YCQ release was sufficient to elicit different responses from the HUVECs. Based on imaging of immunostained cells, the HUVECs that underwent network formation showed about two-fold increase in the expression PECAM-1 when compared to the control samples (**Figure 3D**). PECAM in HUVECs in known to induce enhanced cell mobility and hence network formation. This is a clear indication that the cells were undergoing angiogenic differentiation, in agreement with the results from previous reports.^35,36^

## IV. Conclusion

This study demonstrates the possibility to develop remote-controlled ELMs capable of promoting angiogenic differentiation of vascular endothelial cell cultures in a light regulated manner. This was realized by optogenetically engineering bacteria to produce and secrete a collagen-binding VEGF peptidomimetic fusion protein, YCQ, in response to blue light and securely encapsulating this strain in bilayer hydrogel constructs. With this ELM, we were able to repeatedly switch ON and OFF the release of YCQ up to 9 days and tune the amount released with varying light intensities within the range of 1 – 20 nM in 6 days. The released YCQ was able to adhere to Col-Gel matrices that simulate the ECM of healing wounds and promote network formation in HUVEC cultures, indicating the cells underwent angiogenic differentiation. Notably, the negligible leaky expression of YCQ in dark was insufficient to promote such angiogenic differentiation, thereby validating the capability of controlling the ELM within relevant ranges of YCQ’s functionality.

However, it is desirable to improve the rate and amount of protein released from these ELMs. The low levels of release are expected to be caused by 2 factors – (i) *E. coli* is a poor secretor of proteins^37^and (ii) the PluDA hydrogels have a relatively high polymer content of 30%, possibly resulting in slow protein diffusion.^38^In future studies, both these aspects will be addressed by (i) using gram-positive probiotic bacteria like B. subtilis or lactic acid bacteria, which are prolific at protein secretion^39^ although their genetic toolboxes are poorly equipped compared to *E. coli* and (ii) constructing bilayer hydrogels with lower polymer concentrations of PluDA or introducing soluble fillers that improve matrix porosity.^40^ Despite these limitations, the results in this study clearly highlight the unique advantages that ELMs can offer for regenerative therapies in terms of long-term release of therapeutic proteins and robust in situ control over release profiles.

## V. Materials and methods

### A. Construction of plasmids and bacterial strains

The YebF-CBD-strep-QK fragment (DNA sequence in Supporting information) was ordered as gBlock from Eurofins Scientific, and cloned into pDawn (pDawn was a gift from Andreas Moeglich - Addgene plasmid# 43796; http://n2t.net/addgene:43796; RRID:Addgene_43796) using NEBuilder^®^ HiFi Assembly cloning kit (NEB, E5520S) using following primers-

pDawn Fwd 5’-ataaaagcttAACAAAGCCCGAAAGGAAG-3’,
pDawn Rev 5’-ctctttttttCATGGTATATCTCCTTCTTAAAGTTAAAC-3’,
YebF-CBD-QK Fwd- 5’- atataccatgAAAAAAAGAGGGGCGTTTTTAG-3’
YebF-CBD-QK Rev- 5’- gggctttgttAAGCTTTTATTTCAGGGTC-3’.

This yielded the pDawn-YCQ plasmid which was then transformed into *Clearcoli* BL21(DE3) electrocompetent cells as specified by the provider (BioCat 60810-1-LU). The recombinant pDawn-YCQ was used as a template to construct the pDawn YebF-CBD-Scrambled QK (YCx) mutant (DNA sequence in supporting information) using the NEBuilder^®^ HiFi DNA Assembly Cloning Kit with the following primers:

pDawn Fwd 5’- CTAGCATAACCCCTTGGG-3’,
pDawn Rev 5’- CTAGTAGAGAGCGTTCAC C-3’,
YebF-CBD-Scrambled QK Fwd- 5’- cggtgaacgctctctactagAGTCACACTGGCTCACCTTC-3’
YebF-CBD-QK Scrambled Rev- 5’- gccccaaggggttatgctagTTATTGCTCAGCGGTGGC-3’.

pDawn-YCx was then transformed in *Clearcoli* BL21(DE3) electrocompetent cells as specified by the provider. For storage at −80 °C, glycerol stocks with 30% v/v of glycerol for both clones were made from bacterial cultures grown overnight at 37°C, 250 rpm in the dark from single colonies.

Bacterial culture for protein purification and secretion: 250 mL of *Clearcoli* BL21(DE3) pDawn-YCQ or pDawn-YCx cultures were grown in dark for 37°C, 250rpm in LB Miller medium supplemented with 50 μg/mL of Kanamycin to an OD_600nm_ between 0.4 and 0.8. The culture was then induced for 12 h by exposing it to white light for production of YCQ and YCx at 37 °C, 250rpm

Bacterial cultures for ELMs: *Clearcoli* BL21(DE3) cultures were grown for 16 h at 37 °C, 250 rpm in LB Miller medium supplemented with 50 μg/mL of Kanamycin to an OD_600nm_ around 0.8. All procedures were performed either in the dark or under orange light.

### B. Purification of YCQ and YCx

*Clearcoli* BL21(DE3) with pDawn-YCQ or pDawn YCQx were cultured in LB Miller medium supplemented with Kanamycin. YCQ/YCx production was induced with white light for 16 hours. For harvesting the cells, cultures were transferred to 50 mL flacon tubes and centrifuged with an Avanti J-26S XP centrifuge (Beckman Coulter, Indianapolis, USA) using the JLA-10.500 rotor for 20 min at 4000 rpm and 4 °C. The supernatants were discarded and pellets weighed and stored at −80 °C until further use. For protein extraction, the bacterial pellets were thawed on ice and resuspended in 5 x (cell pellet weight) volume of lysis buffer (100mM TriCl pH8, 150nM NaCl and 1mM PMSF). To lyse the cells, a sonicator (Branson ultrasonics, Gehäuse SFX150) was used with sonication cycles having pulse “ON” for 3 seconds, “OFF” for 5 seconds at 20% power over 6–8 minutes. The sonicated solutions were centrifuged at 14000 rpm for 15 minutes at 4° C and the supernatants were collected for further purification by affinity-based column chromatography. Supernatants and cell debris were stored for analysis by Sodium dodecyl sulfate-polyacrylamide gel electrophoresis (SDS-PAGE). Since the proteins engineered with a StrepII-tag, columns made of Strep tactin beads (Quiagen, 30004) were used for purification of the protein by affinity chromatography modifying the protocol given by the manufacturer(Quiagen). Purification process was optimized to obtain pure proteins. 4ml of strep-tactin bead solution was pipetted into a 15mL falcon tube and was centrifuged at 100rpm for 2 mins to obtain a strep-tactin bed volume of 2mL. The beads were washed three times with 5mL of lysis buffer; the lysis buffer was removed each time by centrifuging the beads at 100rpm for 2 mins. 5ml of cell lysate was added onto the strep tactin beads and this assembly was incubated at 4°C on a rotary shaker for 30 mins to facilitate optimal contact between the beads and cell lysate. The beads were centrifuged at 100 rpm for 2 mins to remove the unbound protein and washed with 5 column volumes of wash buffer(100mM TrisCl, 150mM NaCl and 0.5% tween 20) 3 times by adding 5CV of wash buffer to 15mL falcon and rotating it upside down manually each time; the wash buffer was removed each time by centrifugation parameters mentioned earlier. The protein was eluted using an elution buffer (100mM TrisCl, 150mM NaCl and 2.5mM Desthiobiotin) The eluted protein was rebuffered in 5 volumes PBS pH 7 and concentrated by using 3kD centrifugal filter units and the same centrifugation parameters (4000 rpm for 45 mins at 4°C). The protein yields for YCQ and YCx were in the range of 6 mg/L to 20 mg/L

### C. Patterning of Col-Gel matrix on a Glass Surface to assess binding of YCQ

Coating coverslips with PLL-PEG: Glass coverslips were cleaned by treating them with oxygen plasma in a plasma oven (Harrick Plasma, Ithaca, NY, USA) for 5mins. 50μL of 0.1mg/mL of PLL-PEG (PLL (20)-g [3.5]-PEG (5), SuSoS AG, Dübendorf, Switzerland) solution in PBS was placed on a parafilm; the plasma treated coverslips were inverted on the drop of PLL-PEG. This assembly was incubated for 1h at room temperature (RT).

i. Cleaning of the photomask: The patterned surface of the photomask was cleaned with acetone, water, ethanol and water respectively. It was air dried gently. It was placed in UVO cleaner for 5 min, in such a way that the patterned side faces up.
ii. Making patterns with the help of photomask: After the incubation, the coverslip was released from the parafilm gently by infusing deionised water under the coverslip in such a way that it starts floating. It was rinsed by dipping the patterned surface 5 times in PBS pH 7 solution and dried by draining the access solution onto a paper towel. 5μL of deionised water was placed on the patterned side of the photomask in order to reduce friction between the coverslip and patterned surface. The coverslip was carefully picked up with forceps and placed on the drop of water on the patterned surface. This assembly was placed in UVO cleaner (Jelight Company Inc, model no.42) for 5 mins in a way that the pattered surface holding coverslip faces down. UV exposure cleaves PEG chains at the site of exposure on the glass coverslip. The coverslip was released gently by infusing 100 μL PBS pH 7beneath it and it was washed with PBS by dipping it in PBS 5 times. The coverslip was then placed on the 25 μL Col-Gel solution which was prepared by modifying a reported protocol.^32^ All stock solutions were made in PBS pH 7. The gels were prepared by mixing 75% v/v of 2mg/ml collagen (gibco, A1048301) and 25% v/v of 2mg/ml gelatin (SigmaAldrich, 9000-70-8) solution. The two solutions were mixed by manually rotating the pipette tip in the Eppendorf tube without pipetting it. The gels were allowed to polymerize at 37 °C, 5% CO2 for 1 hour
iii. Incubation of YCQ and YCx supernatants or purified solutions and staining on the patterned surface: Supernatants from overnight induced (light) and uninduced (dark) YCQ and YCx were collected by centrifuging the bacterial culture at 4000rpm for 20 mins. The supernatants were filter sterilized using 0.4 μm syringe filters (Carl roth, SE2M230104). The coverslips with Col-Gel patterns were then incubated with either filtered supernatants or 10 nM purified solutions of YCQ and YCx by inverting the coverslips on 25 μL drops of protein solutions. This assembly was incubated for one hour at RT. After the incubation, glass coverslips were released gently from the photomask by infusing 100μL of PBS between glass surface and the parafilm. The coverslips were picked up with forceps and dipped in PBS 5 times to remove the unbound protein. The patterned glass surfaces with immobilized proteins were then incubated with anti-YebF primary antibody (Athen ES, AES-0313) by placing the patterned coverslip on 25μL of the 1:500 diluted (in 1% BSA) antibody on parafilm. After incubation at RT for one hour the coverslips were released as mentioned before and all the patterned surfaces were then incubated with AF-488 Goat anti rabbit secondary antibody (ThermoFischer, A-11008) for one hour at RT. Patterned surfaces were washed with PBS as mentioned before and were placed on a glass slide by placing 10uL of PBS on the glass slide and inverting the patterned surface on it. YCQ or YCx stained Col-Gel patterns were visualised using a Nikon Ti-Eclipse microscope (Nikon Instruments Europe B.V., Germany).

### D. Fabrication of ELM

i. <u>Silanizing glass coverslip</u>: Glass coverslips (13mm) were arranged in a Teflon holder with a removable handle (custom made) for washing steps in a beaker. The teflon holder with coverslips was suspended in a beaker with 99% ethanol and was sonicated for 10 minutes. Then, it was washed with ultrapure water followed by 99% ethanol. The Teflon holder handle was removed and the base holding the cover slips was slid into a 50 mL falcon tube containing 20 mL of 95% ethanol, 4% milli Q water and 1% of 3 – APS (3-Trimethoxy silyl) propyl acrylate (Sigma-Aldrich, 4369-14-6) such that coverslips were completely immersed in the solution. After incubation overnight at RT, the coverslips were washed 3 times with deionised water to remove access APS and transferred into a beaker with ultrapure water and stored in it till further use.
ii. <u>Preparation of Polydimethylsiloxane (PDMS) moulds</u>: A beaker in which PDMS moulds were to be made was sonicated for 3 minutes with 99 % ethanol in it._10 grams of SYLGARD™ 184 silicone elastomer base (Dow chemicals, USA) was added to beaker on a weighing balance and 1 gram of SYLGARD™ 184 silicone elastomer curing agent (cross linker) (Dow chemicals, USA, 1023993) was also added and mixed thoroughly using a spatula. The Beaker with the PDMS-crosslinker mixture was placed in a vacuum-desiccator (DN 150 Duran) up to 10 minutes to remove air bubbles from the mixture. The beaker was then removed from the desiccator after confirmation of absence of bubbles in mixture. The volume of the PDMS mixture was calculated to obtain moulds of dimensions depicted in Figure 2 and Figure S4 and poured into a glass petri dishes. These were then placed in a hot air oven at 95 °C for 2 hours for polymerization. They were taken out after 2 hours and left undisturbed at room temperature overnight._Solidified PDMS moulds were scraped out from glass flasks and holes of desired diameter were punched with wad punch set (BOEHM, 832100).
iii. Preparation of Pluronic diacrylate solution: Pluronic diacrylate (PLU-DA) was obtained from the group of Prof. del Campo, who synthesized it as previously reported.^41^ 30% Pluronic diacrylate (PLU-DA) + 0.02 % IRGACURE solution was prepared for construct fabrication. 3 grams of PLU-DA and 20 mg of IRGACURE 2959 (2-Hydroxy-4’-(2-hydroxyethoxy)-2-methylpropiophenone) (Sigma Aldrich, 410896-10G) photo initiator were dissolved in ultrapure water in an amber coloured glass bottle and the final volume was adjusted to 10 mL. The PLU-DA solution was kept on a rotary shaker at 4 °C overnight for complete dissolution of contents. PLU-DA was stored in the same amber glass bottle at 4 °C until further use.
iv. Fabrication of ELMs: 5 mL bacterial cultures at their exponential phase (OD = 0.5-0.8) were centrifuged at 4000 rpm for 10 minutes at room temperature. Pellets were resuspended in M199 cell culture media and was made up to an OD of 1 after measuring it in a Nanodrop One device. Bacterial suspensions were prepared by mixing 90% volume of PLU-DA and 10% volume of bacterial culture (OD = 1) such that the final density of bacteria in PLU-DA mixture was 0.1 OD_600nm_. Solutions containing PLU-DA were always handled on ice to prevent their physical gelation. 3-APS coated coverslips were placed on disinfected paraffin paper in a petri dish. A cylindrical PDMS mould with 6 mm diameter was placed on the coverslips with conformal contact. 22 μL of bacterial gel solutions were pipetted into the PDMS moulds to make a core layer of radius 3 mm and height of 0.7 mm. It was left undisturbed for 2 minutes at room temperature for the PLU-DA to form a physical gel. The PDMS + coverslip setup was transferred to a Gel-Doc (Biozym Scientific GmbH, FluorChemQ) and the PLU-DA-Bacterial mixture was cross-linked by irradiating it with low intensity UV transillumination (365 nm) for a duration of 60 seconds. The whole setup was transferred back to a sterile hood and left undisturbed for 2 minutes after which the PDMS mould was removed from the coverslip. A PDMS mould of 10 mm diameter was placed on the same coverslip to produce a protective PLU-DA shell to prevent the leakage of bacteria from the ELM. 60 μL of PLUDA was pipetted onto the PLU-DA bacterial core layer such that it formed a shell layer covering the entire surface of core with a diameter of 10 mm and height of 1 mm. The PDMS and coverslip setup was transferred to the Gel-Doc and cross-linked by UV transillumination for a duration of 90 seconds. The whole setup was transferred back to the sterile hood and the PDMS mould was removed from the coverslip. This ELM construct was transferred into a well in a 24-well plate and 300 μL of M199 cell culture media supplemented with additional 0.4% glucose and 0.88% NaCl was pipetted into the well. The 24-well plates with hydrogel constructs was placed in an incubator (28 °C) and after 16 hours in dark. For reversible light switching experiments.: one 24-well plate was kept in light [blue light irradiation device; pulse: 2 sec ON, 1min OFF;200 μW/ cm^2^] for 3 days and then moved to dark for 3 days followed by a period of light again for 3 days. The other 24 well plate was kept in dark for 3 days and then switched to light (same irradiation parameters as mentioned before) for 3 days followed by a period of dark for 3 days. Surrounding media was collected from each well after every 3 days and ELMs were replenished with 300μL of fresh media. For light tuning experiments: 24 well plate containing ELMs was kept on opto well irradiation device which was programmed to irradiate wells with blue light pulses (2sec ON; 1 min OFF, 450nm) of intensities of 0μW/cm^2^, 80 μW/cm^2^, 105 μW/cm^2^ and 125 μW/cm^2^ for 6 days. The volume of media surrounding ELM was decreased in 3 days from 300 μL to 250 μL due to evaporation. It was replenished by adding 50μL of the fresh media to it. The surrounding media containing secreted protein was collected at day 6 for quantification with ELISA (see point vi)
v. Qdot Streptavidin staining of ELMs: ELMs in 24-well plate wells were washed with PBS thrice. Qdot 655 Streptavidin conjugate (Thermo Fisher scientific, Germany, Q10121MP) was used to stain the hydrogels. The staining solution was prepared in the ratio of 1:500 in 2% BSA. 300 μL of staining solution was added to each ELM construct and was incubated at 37 °C for 2 hours. The well plate was completely wrapped with aluminium foil to avoid photobleaching during incubation. The antibody solution was pipetted out from wells and constructs were washed with 300 μL of PBS thrice. Then constructs were analysed by Nikon epifluorescence microscope and overlay images of both brightfield as well as the 561 nm red channel were captured.
vi. Analysis of ELM supernatant by ELISA: ELISA was performed using Strep-tactin coated 96 well plates (iba-life sciences) with the following protocol:

1. An initial blocking step was performed with 100 μL of 2% Bovine serum albumin (BSA) added to the wells and the well-plate left overnight at 4 °C or for 1 hour at 37 °C.
2. The wells were then washed by pipetted out the blocking solution and washing with 100 μL of wash buffer (PBS + 0.1% Tween-20) thrice.
3. 100 μL of samples were added to the wells and left to incubate for 1 hour at room temperature followed by 3x washing step.
4. Another blocking step was performed where 100 μL of 2% BSA was added to wells and left for 1 hour at room temperature followed by 3x washing step.
5. Primary antibody treatment: Rabbit Anti-YebF antibody (Athena enzyme systems, USA) was prepared in the ratio of 1:500 in 2% BSA. 100 μL of antibody solution was added to wells and left at room temperature for 1 hour followed by 3x washing step.
6. Secondary antibody treatment: Goat anti-Rabbit IgG conjugated with horseradish peroxidase (Invitrogen) was used as secondary antibody in the ratio of 1:500 in 1% BSA. 100 μL of secondary antibody solution was added to wells and left at room temperature for 1 hour followed by 4x washing step.
7. Substrate treatment: 100 μL of Tetramethylbenzidine (TMB) (Sigma-Aldrich, Germany) substrate was added to each well and was left undisturbed until stable blue color developed.
8. Stopping step: 100 μL of 1M Hydrochloric acid (HCl) was added to wells to stop the reaction and it resulted in yellow coloration of solution which was measured within 30 minutes.
9. The absorbance of test samples was measured at 450 nm by using *iControl 2000* software of Tecan plate reader and resulting data was analysed.

Using serial dilutions of the purified proteins in the optimized M199 medium, a standard curve was plotted. The slope of this standard curve enabled determination of the molar concentrations of ELM-released YCQ or YCx.

### E. HUVEC network formation assay

i. Cell Culture Conditions: HUVECs (PromoCell, C-12205) were maintained on cell culture flasks coated with gelatin (0.2%). Cells were cultured in M-199 medium (Sigma, M4530) supplemented with penicillin (1000 UL-1), streptomycin (100 mgL-1, Sigma), ECGS (Sigma, E2759), sodium heparin (Sigma, H-3393), and 20% fetal bovine serum (FBS, Gibco, 10270) as previously described^35^HUVECs between passages 2 to 7 were used for the experiments.
ii. Preparation of Col-Gel surfaces: Col-Gel surfaces were prepared as mentioned previously (Materials and Methods section C ii) 10uL of this solution was added into the 15 well angiogenesis well plates (ibidi, 81506). The gels were allowed to polymerize at 37 °C, 5% CO2 for 1 hour followed by overnight incubation at 4°C.
iii. Immobilization of secreted YCQ on Col-Gel surfaces: 10 μL of the SN collected from ELMs activated with light for 6 days were incubated on the Col-Gel surface for 1 hour at 37°C in the CO2 incubator. The supernatants were removed with a pipette and the Col-Gel surfaces were washed with 50 μL of PBS once to remove inbound proteins.
iv. HUVEC seeding: 50 μL of 2×10^5^ cells/mL HUVECs (P3-P7) were seeded on the Col-Gel surfaces and the plate was incubated at 37°C and 5% CO_2_ for 16 hours. The culture was then fixed with 4% aqueous PFA solution for 15 mins, washed with PBS and blocked with 5% BSA solution for 1 h. Cells were permeabilized with 0.5% Triton X-100 for 15 h and incubated with monoclonal goat anti-rabbit PECAM-1 primary antibody (1:500 in 1% BSA for, Abcam) overnight and washed with PBS. This was followed by incubation with anti-rabbit Alexa flour-488 secondary antibody (1:500 in water, Thermo Fisher Scientific) for 1 h. Subsequently, cells were washed with PBS and nuclei were stained with DAPI (1:500 in water, Life Technology) and actin fibers were labelled with TRITC-phalloidin (1:500 in water, Thermo Fisher Scientific). Samples were washed thrice with PBS and imaged with a Nikon epifluorescence microscope at 10X magnification using excitation wavelengths of 405 (900ms excitation), 488(100ms excitation), and 565(100ms excitation) with the same excitation duration for each channel and 20% incident light intensity. Images were captured using NIS-Elements software and processed as follows for inclusion in the figures: Image processing and analysis was done using Fiji edition of ImageJ (Image J Java 1.8.0). Brightness and contrast for the DAPI(Cyan, 405) and Actin (Yellow, 488) were adjusted in their respective LUT histograms to the same range for all the images in an individual experiment. For PECAM-1, since Anti-PECAM-1 agglomeration at some places created bright spots to different degrees, the brightness and contrast in the LUT histograms were manually adjusted for each image based on the maximum intensity within the cells that were not caused by the agglomerates and the minimum intensity in the cell-free background. This way ensured best possible visualization of PECAM-1 expression in all the cells.
v. PECAM-1 analysis - Fiji edition of ImageJ (ImageJ Java 1.8.0). was used for quantification PECAM-1 fluorescence intensity. Quantification of the fluorescence intensities was done by determining the mean grey value within the cells excluding the bright spots created by agglomerates (by manual selection) and subtracting the mean grey value of the cell-free background.

#### Statistical analysis

The data are expressed as the individual data points ± standard deviation and presented without further pre-processing. Each data point represents mean value of an individual sample. All experiments comprised of at least two independent experimental batches performed under identical conditions.

## Acknowledgments

The authors acknowledge support from the Leibniz Science Campus on Living Therapeutic Materials, LifeMat and the Deutsche Forschungsgemeinschaft’s Collaborative Research Centre, SFB 1027. pDawn was a gift from Andreas Moeglich (Addgene plasmid# 43796; http://n2t.net/addgene:43796; RRID:Addgene_43796). The authors thank Shardul Bhusari, Hafiz Syed Usama Bin Farrukh and Samuel Pearson (INM – Leibniz Institute for New Materials) for chemical synthesis of Pluronic F127 diacrylate and Claudia Fink-Straube (INM – Leibniz Institute for New Materials) and Klaus Hollemeyer (Saarland University) for performing the MALDI-TOF measurements.

