## Supporting information for "Light-regulated pro-angiogenic engineered living materials"

**Gene sequence of YCQ : YebF-CBD-Streptag-QK**

GTTTAACTTTAAGAAGGAGATATACCATGAAAAAAAGAGGGGCGTTTTTAGGGCTGTTGTTGGTTTCTG
CCTGCGCATCAGTTTTCGCTGCCAATAATGAAACCAGCAAGTCGGTCACTTTCCCAAAGTGTGAAGATCT
GGATGCTGCCGGAATTGCCGCGAGCGTAAAACGTGATTATCAACAAAATCGCGTGGCGCGTTGGGCAG
ATGATCAAAAAATTGTCGGTCAGGCCGATCCCGTGGCTTGGGTCAGTTTGCAGGACATTCAGGGTAAAG
ATGATAAATGGTCAGTACCGCTAGCCGTGCGTGGTAAAAGTGCCGATATTCATTACCAGGTCAGCGTGG
ACTGCAAAGCGGGAATGGCGGAATATCAGCGGCGTGGTACCACTGGAGGATGGCGCGAACCGAGCTTT
ATGGTGCTGAGCGGCGGCGGTTCGGGGTCGGGATCGGGATCATGGAGCCACCCGCAATTTGAGAAAG
GCTCGGGCTCGGGGAGTGGAAGTGATATTGGCAAATATAAACTGCAGTATCTGGAACAGTGGACCCTG
AAATAAAAGCTTAACAAA

Gene sequence of YCx: **YebF**-**CBD**-**Streptag**-QK

AAGGAGATATACCATGAAAAAAAGAGGGGCGTTTTTAGGGCTGTTGTTGGTTTCTGCCTGCGCATCAGT
TTTCGCTGCCAATAATGAAACCAGCAAGTCGGTCACTTTCCCAAAGTGTGAAGATCTGGATGCTGCCGG
AATTGCCGCGAGCGTAAAACGTGATTATCAACAAAATCGCGTGGCGCGTTGGGCAGATGATCAAAAAA
TTGTCGGTCAGGCCGATCCCGTGGCTTGGGTCAGTTTGCAGGACATTCAGGGTAAAGATGATAAATGGT
CAGTACCGCTAGCCGTGCGTGGTAAAAGTGCCGATATTCATTACCAGGTCAGCGTGGACTGCAAAGCGG
GAATGGCGGAATATCAGCGGCGTGGTACCACTGGAGGATGGCGCGAACCGAGCTTTATGGTGCTGAGC
GGCGGCGGTAGTGGTAGTGGTAGTGGTAGTTGGAGCCACCCGCAATTTGAGAAAGGCTCGGGCTCGG
GGAGTGGAAGTGGCTTACGTCATAAGCGTCCGTCACAAGAGAAATTGGGTTAAAAGCTTAAC

**SDS-PAGE and Western Blot**: YCx and YCQ proteins were purified from 250 mL cultures as mentioned in the Materials and Methods section and 5 µL Laemmli 4x buffer was mixed with 15uL of the both the proteins and heated at 95°C for 5 min. 10 µl of this was loaded on separate 15% SDS gels and run at 120 V for 60 min. YCx gel was Biosafe Coomassie (Bio-Rad) stained. YCQ gel was wet blotted to a PVDF membrane (Merck Millipore) with Mini Trans–Blot Cell (Bio-Rad) at 100V for 1h. Qdot streptavidin (Thermo Fisher scientific, Germany, Q10121MP) staining was performed with SNAP i.d. 2.0 (Millipore). diluted 1:500 in PBST. Pictures were taken in RGB multiplex mode of Fluorchem Q imager.

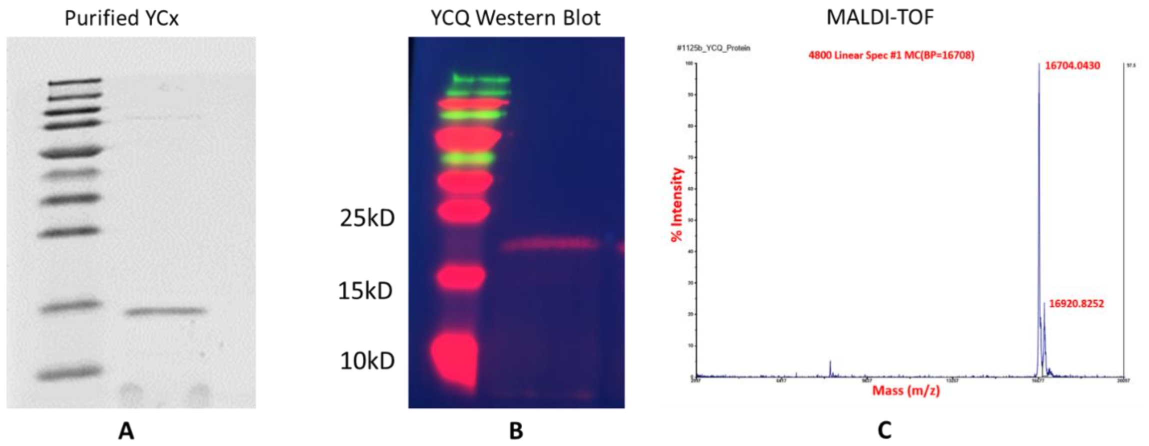


**Figure S1:** A. Purified YCx on 15% Polyacrylamide gel B. Western Blot for confirmation of YCQ. YCQ was stained with fluorescently labelled Q dot streptavidin (red) C. MALDI-TOF MS spectrum of purified YCQ protein; sharp peak at 16.704 kDa confirms the size of YCQ protein after sec-leader peptide cleavage.

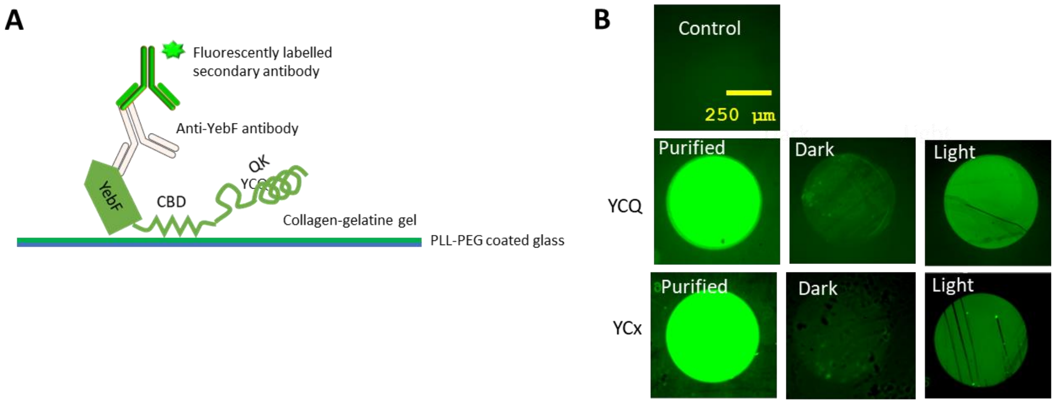


**Figure S2**: YCQ adhesion to Col-Gel Matrices A. Graphical representation of YCQ and YCx adhesion to Col-Gel matrices. B. YCQ and YCx was (purified and 12h culture SNs) immobilized on Col-Gel matrices and were stained with 1:500 diluted rabbit anti YebF Primary antibody followed by staining with 1:500 diluted goat anti-rabbit AF-488. The samples were imaged in Nikon epifluorescence microscope using 10X objective; Excitation; 488

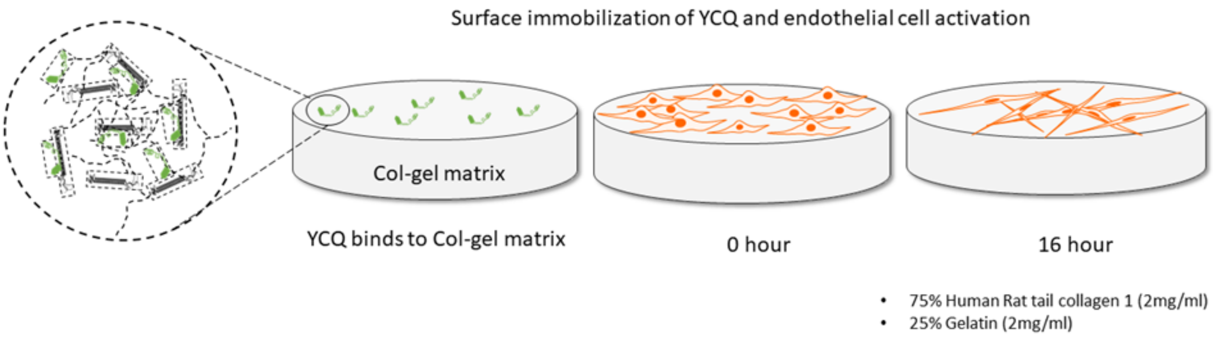


**Figure S2**: Col-Gel wound healing model for promotion of angiogenesis in HUVECs. YCQ is immobilized on the surface of col-gel gels. HUVECs undergo morphological change and from cell networks within 16 hours of seeding the on the col-gel YCQ gels

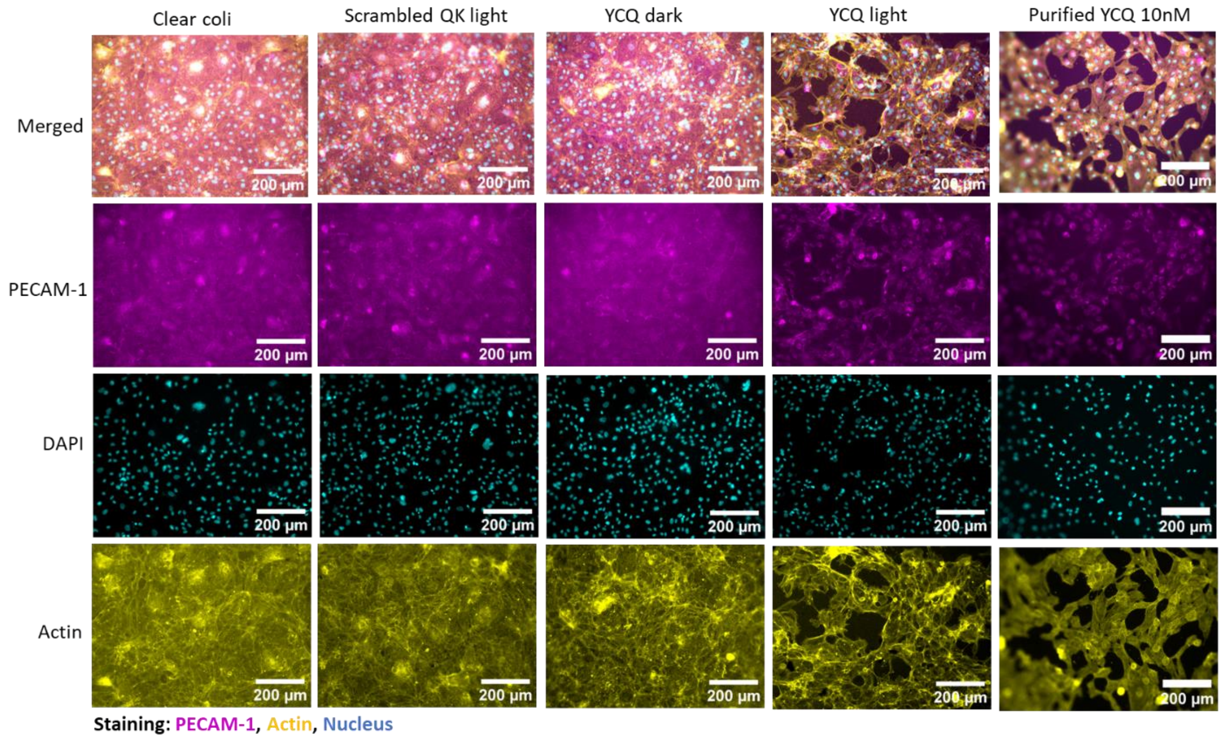


**Figure *S*4:** Epifluorescence images of HUVECs after 16 hours of incubation on col-gel surface functionalized with Clearcoli (control), YCx (negative control), and YCQ SNs from liquid culture after 12 hours of induction with blue light. HUVECs are labelled with PECAM-1 antibody (magenta), DAPI (cyan) to stain nucleus and Phalloidin-AF488 (yellow) to stain actin fibres).

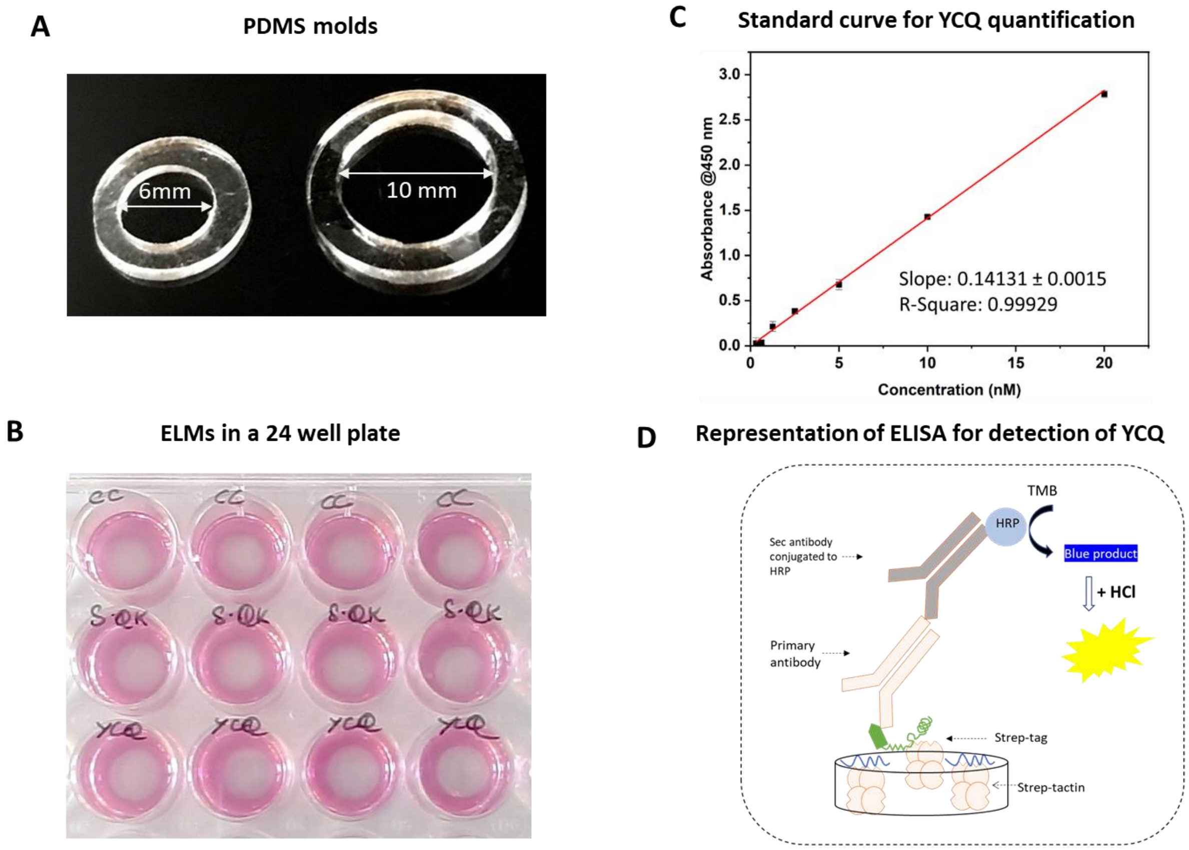


**Figure S5**: A. PDMS molds for ELM fabrication B. ELMs in a 24 well plate suspended in growth supporting media C. ELISA standard curve for YCQ quantification D. Representation of ELISA for quantification of YCQ. YCQ was immobilized on the surface of strep-tactin coated 96 well plates and stained with rabbit anti-YebF primary antibody followed by AF-488 goat anti rabbit secondary antibody.

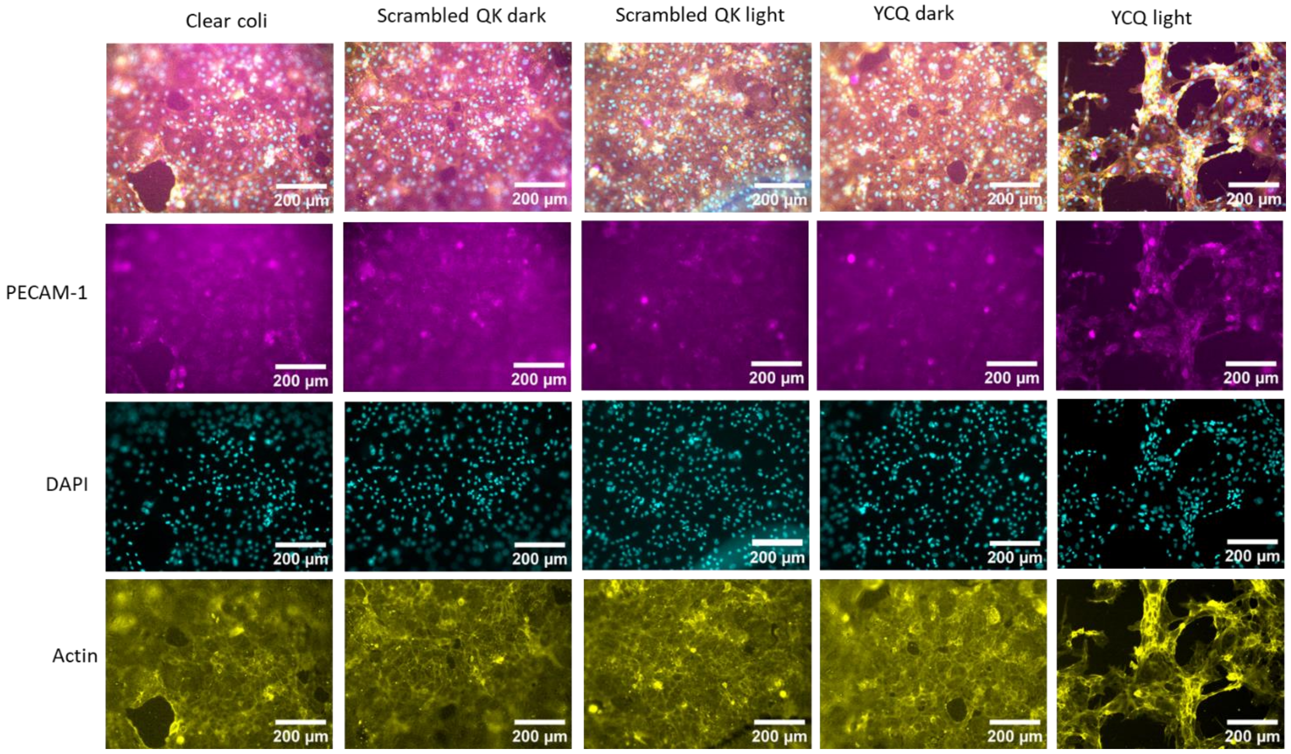


**Figure S6:** Replicate 1: Epifluorescence images of HUVECs cultured for 16 hours on Col-Gel surfaces that were incubated in supernatants from ELMs containing unmodified bacteria (control), YCx (negative control), and YCQ kept in dark or exposed to blue light pulses (2sec ON, 1min OFF; 105 µW/cm2) for 6 days. The cells were stained for PECAM-1 (magenta), actin (yellow) and DNA (cyan).

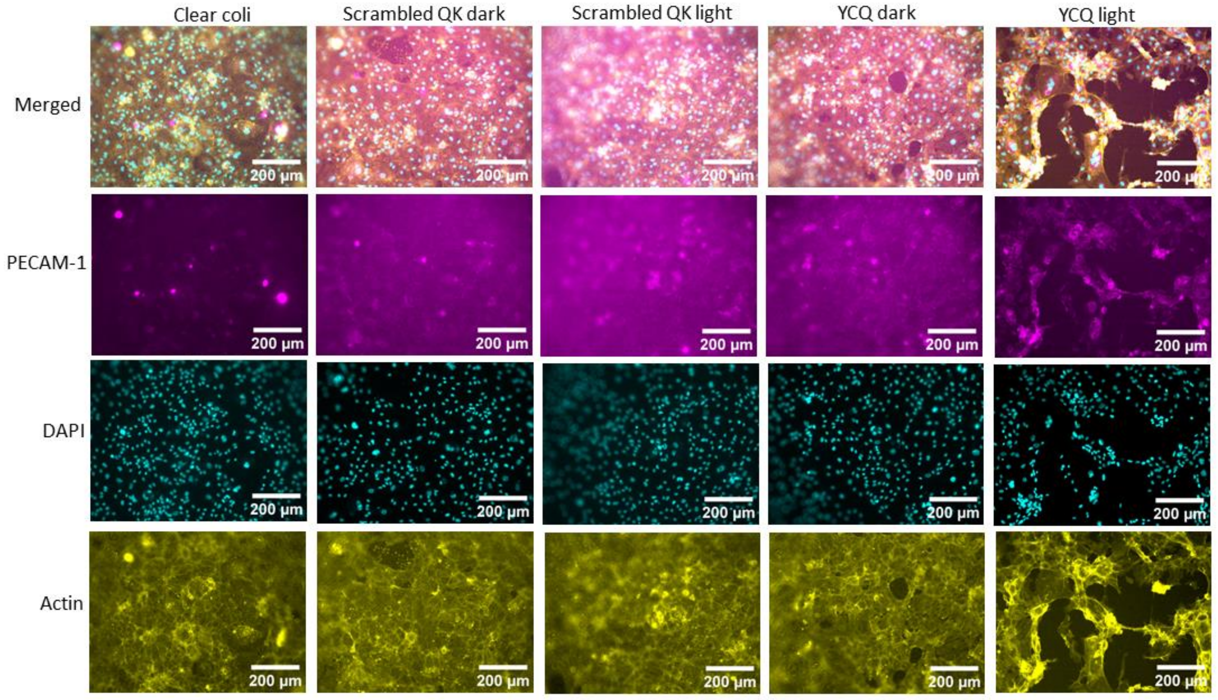

**Figure S7**: Replicate 2: Epifluorescence images of HUVECs cultured for 16 hours on Col-Gel surfaces that were incubated in supernatants from ELMs containing unmodified bacteria (control), YCx (negative control), and YCQ kept in dark or exposed to blue light pulses (2sec ON, 1min OFF; 105 µW/cm2) for 6 days. The cells were stained for PECAM-1 (magenta), actin (yellow) and DNA (cyan).

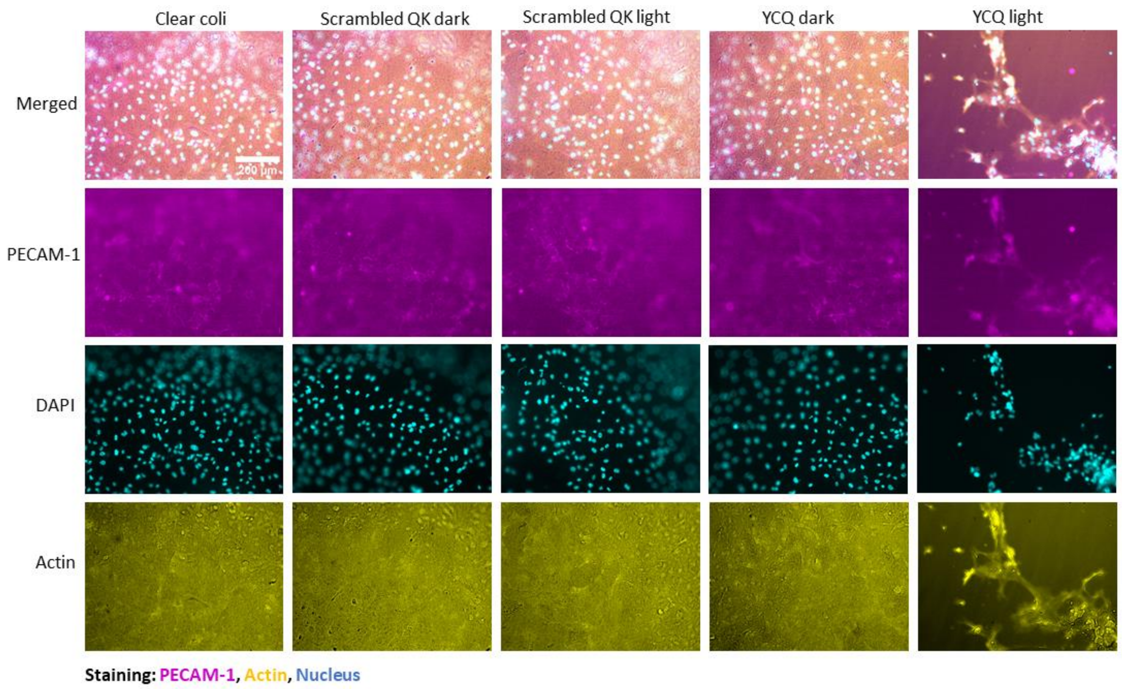


**Figure S8**: Replicate 3: Epifluorescence images of HUVECs cultured for 16 hours on Col-Gel surfaces that were incubated in supernatants from ELMs containing unmodified bacteria (control), YCx (negative control), and YCQ kept in dark or exposed to blue light pulses (2sec ON, 1min OFF; 105 µW/cm2) for 6 days. The cells were stained for PECAM-1 (magenta), actin (yellow) and DNA (cyan).
